# Humanized FLT3 mice display enhanced tissue engraftment and support HIV‑1 persistence and rebound

**DOI:** 10.64898/2026.06.03.727073

**Authors:** Patricia Resa-Infante, Patricia Piñol, Fernando Laguía, Gerard Campos-Gonzalez, Àlex Cobos, Victor Urrea, Maria del Carmen Garcia-Guerrero, Yaiza Rosales-Salgado, Marina Cid, Jorge Díaz-Pedroza, Leonard D Shultz, Voot Yin, Michael A. Brehm, Brian Soper, María Salgado, Javier Martinez-Picado

## Abstract

Despite advances in antiretroviral therapy (ART), HIV-1 cure efforts remain hindered by viral reservoirs in long-lived myeloid cells and immune-privileged tissues that are less accessible and therefore unlikely to be assessed in human clinical trials. Consequently, there is a critical need for robust research platforms such as immune cell humanized mice to bridge preclinical and clinical HIV research. However, previously described humanized mouse models have demonstrated incomplete hematopoietic development, particularly showing low levels of NK or myeloid cells. Herein, we present the novel humanized FLT3 mouse model that develops NK cells, myeloid progenitors, monocytes, and both functional conventional (cDCs) and plasmacytoid dendritic cells (pDCs) to support HIV-1 infection.

Human cord blood derived CD34^+^ hematopoietic stem cells (HSC) were engrafted in the FLT3 (Hu-FLT3) and NSG (Hu-NSG) mouse strains for comparison. Our data showed that while Hu-NSG and Hu-FLT3 mice have comparable human lymphocyte levels, the proportion of myeloid cells (including monocytes, pDCs and cDCs) in Hu-FLT3 mice (16.2 %) was three-fold higher than in Hu-NSG mice (5.6 %) and the proportion of NK cells was six-fold higher (12.8 % and 1.9 %, respectively). Both strains successfully supported HIV-1 infection, maintain viral replication for 17 weeks in untreated mice, and proviral DNA was detectable in peripheral blood, bone marrow and spleen. While ART effectively reduced viral load to undetectable levels in four weeks in both strains, we observed viral rebound after treatment discontinuation within 3 weeks, reaching the same levels of viral load pre-ART and mimicking what is observed in people living with HIV (PLWH).Human immune cells and HIV-1 RNA were higher in tissues of Hu-FLT3 mice compared to Hu-NSG mice, mirroring features reported in human tissue reservoirs.

Our findings demonstrated that Hu-FLT3 mice support enhanced development of human innate immune cells in blood and tissues, which are associated with higher levels of HIV-1 replication compared to Hu-NSG mice. This study establishes a novel, robust and accessible *in vivo* platform to investigate potential HIV cure and persistence-targeting interventions with translational relevance to human therapeutic development thanks to the improved and more complete human immune repertoire in Hu-FLT3.

## 1. Introduction

For people living with HIV, lifelong antiretroviral therapy (ART) remains the cornerstone of treatment effectively suppressing viral replication and managing the progression of the disease. The major obstacle to achieving a definitive cure is the persistence of viral reservoirs in blood cells and tissue compartments, particularly in resting memory CD4^+^ T cells and myeloid cells (including monocytes, macrophages, microglia, and dendritic cells) (1–7). Myeloid cells have been understudied in the context of HIV cure because their proportion in the human body is lower compared to other immune populations and they are mainly present in less accessible tissues. Thus, there has been limited study of the role of myeloid cells in regular and non-invasive clinical trials (8). Only recent studies, with greater tissue sampling at the end of life, help us to understand the nature of viral reservoir in all the tissues (9).

Consequently, there is a critical need for robust research platforms to facilitate the preclinical evaluation of novel antiviral drugs and vaccine candidates. Although non-human primate models provide important translational insights, their use is constrained by cost, experimental duration, specialized infrastructure, and ethical considerations (10). Small animal models, particularly humanized mice, offer a complementary and more accessible *in vivo* platform for mechanistic studies and intervention testing. Currently available humanized mouse models used in HIV cure studies include: peripheral blood mononuclear cell humanized mice (Hu-PBMC), hematopoietic stem cell humanized mice (Hu-HSC), and bone marrow-liver-thymus humanized mice (Hu-BLT) generated by the implantation of human fetal liver and thymus tissues together with injection of HSC (11), each with distinct advantages and limitations such as imperfect immune reconstitution and inadequate lymphoid architecture (12). The Hu-BLT model provides the most complete human immune repertoire reconstitution, but the limited fetal sample source poses a major restriction, together with complicated logistics and ethical constraints in different countries (13). On the contrary, Hu-PBMC and Hu-HSC models do not require complex facilities and several of these models are commercially available, which facilitates their use in laboratories without experience in humanization. Hu-PBMC-based models that allow rapid engraftment are restricted by graft-versus-host disease, limiting their use to short-term studies (14). Hu-HSC models support long-term studies such as antiviral treatments and/or HIV latency establishment experiments but typically exhibit incomplete immune reconstitution and do not mimic full human hematopoietic repertoire.

The number of immunodeficient murine strains available for humanization has increased significantly during the last several years by genetic engineering and the NOD *scid* gamma strain (NSG, NOD.Cg-*Prkdc*^*scid*^ *Il2rg*^*tm1Wjl*^/SzJ) has become the most frequently used strain (15). NSG mice are deficient in mature host lymphocytes and NK cells and survive for long periods of time without lymphoma development. The lack of IL2 receptor gamma chain in NSG mice has supported heightened engraftment compared to previous NOD-scid strains (16). Hence, NSG mice transplanted with human cord blood-derived CD34^+^ HSC (Hu-NSG) are the current gold standard for immunology, immuno-oncology, and infectious disease research that require *in vivo* models with human immune system reconstitution. We and others have largely used Hu-NSG mice with good results (17,18), although this model has limited development of human NK and myeloid lineages. An alternative to NSG is the triple transgenic NSG-SGM3 (NSGS) mice, expressing human stem cell factor (SCF), granulocyte-macrophage colony-stimulating factor (GM-CSF) and interleukin-3 (IL-3) (19). It combines the features of the highly immunodeficient NSG mouse with expression of human cytokines that support the stable engraftment of myeloid lineages and regulatory CD4^+^ T cell populations despite limited NK maturation. Another variant is the transgenic NOD.Cg-*Prkdc*^*scid*^ *Il2rg*^*tm1Wjl*^ Tg(IL15)1Sz/SzJ (Hu-IL15) model where human IL15 expression enhances human NK cell development upon human stem cell engraftment, but human myeloid lineage development remains impaired (20).

A recently developed mouse strain in the NSG background has been genetically modified to knock out the endogenous mouse FLT3 receptor and knock in a human transgene containing the FLT3 ligand (21,22), expanding the single elimination of this receptor (23). This mouse strain, NOD.Cg-*Flt3*^*em2Mvw*^ *Prkdc*^*scid*^ *Il2rg*^*tm1Wjl*^ Tg(FLT3LG)7Sz/SzJ, when engrafted with human CD34^+^ HSC (Hu-FLT3) improved differentiation of monocytes and dendritic cell subsets such as CD33^+^ myeloid progenitors, CD14^+^ monocytes, and both functional CD11c^+^ conventional and CD123^+^ plasmacytoid dendritic cells (cDCs and pDCs, respectively), in addition to NK cell development. These features pose an added advantage of this humanized mouse model as a preclinical platform to investigate infectious diseases with strong tissue and innate immune involvement.

In this study, we have characterized the novel Hu-FLT3 strain in the context of HIV-1 infection in direct comparison to the Hu-NSG standard model. We evaluated Hu-FLT3 mice in terms of i) human immune reconstitution in blood and tissues; ii) susceptibility to HIV-1 long-term infection; iii) response to oral antiretroviral therapy (ART) to reduce animal manipulation; iv) viral rebound following analytical treatment interruption; and v) HIV-1 persistence in blood and lymphoid organs. In addition, we assessed the robustness and translational relevance of the Hu-FLT3 model for preclinical HIV research by incorporating donor-to-donor variability.

## 2. Material and methods

### 2.1. Ethics and biosafety statements

All animal experiments using infectious HIV-1 were performed in the BSL3 laboratory of the Center for Bioimaging and Comparative Medicine (CMCiB-IGTP). Mice were housed in ventilated cages at a temperature of 18°C–22°C, humidity of 50–70%, 50 air exchanges per hour in the cages, a 12/12-h light/dark cycle and with five mice as maximum cage density. Mice were fed a standardized pellet diet (unless ART was administered, method details below) and provided sterilized water *ad libitum*. All materials were autoclaved before use. Animals were supervised daily following a strict protocol to ensure animal welfare. The study was approved by the institutional review board for biomedical research of Hospital Germans Trias i Pujol (Ref. PI-24-201). All animal procedures described in protocol 12098 were reviewed by the Animal Experimentation Ethics Committees from the same hospital and approved by Generalitat de Catalunya, according to current Spanish and European Union legislation regarding the protection of experimental animals.

### 2.2. Humanization of FLT3 and NSG mice

NOD.Cg-*Flt3*^*em2Mvw*^ *Prkdc*^*scid*^ *Il2rg*^*tm1Wjl*^ *Tg(FLT3LG)7Sz*/SzJ (FLT3, strain 033367) and parental NOD.Cg-*Prkdc*^*scid*^ *Il2rg*^*tm1Wjl*^/SzJ (NSG, strain 005557) mice were generated at The Jackson Laboratory (USA). Four-week-old females were sub-lethally irradiated (140cGy) and injected via tail vein with 5-6 × 10^4^ umbilical cord blood CD34^+^ HSC from human female or male donors. Female recipients were exclusively used due to greater engraftment efficiency and universal acceptance of both male and female donor cells. Mice were bred and housed at The Jackson Laboratory until the validation of hematopoietic reconstitution at 12 weeks post-transplant by flow cytometry using a panel of antibodies against the following markers: hCD45, hCD19, hCD3, CD33, CD56. Animals with more than 25% of human CD45^+^ cells in the circulation were transported to the BSL3 laboratory of the CMCiB-IGTP animal facility for experimental performance.

In this study, 30 FLT3 mice humanized with CD34^+^ HSC from seven different donors (n = 4-7) and 14 NSG mice humanized with CD34^+^ HSC from two donors (n = 7 each) for comparison were randomly allocated into three experimental groups. Engraftment data was collected prior to infection between week 15 and week 19 post-transplant. Uninfected mice (Ctrl group) received PBS and were followed until the study endpoint (Hu-FLT3 from two donors n = 4; Hu-NSG from two donors n = 4). The rest of the animals were intravenously infected with HIV-1_NL4-3_ virus to monitor viral replication for 5 weeks (Hu-FLT3 from five donors n = 26; Hu-NSG from two donors n = 10). A subgroup of these mice was monitored for 17 weeks to evaluate long-term infection (Hu-FLT3 from two donors n = 4; Hu-NSG from two donors n = 4) and compared to a parallel subgroup that at week 5 post-infection initiated oral ART for 6 weeks, followed by analytical treatment interruption (ATI) for 6 weeks to assess viral rebound (Hu-FLT3 from two donors n = 6; Hu-NSG from two donors n = 6). Sample sizes varied by availability of samples/timepoints.

### 2.3. Immunophenotyping of blood cells in engrafted mice

Submandibular bleeds were performed to collect 40–100 µL of blood from mice in EDTA 2ml-tubes. Plasma was stored separately upon centrifugation for 5 minutes at 800 x g, and cells were incubated with fresh erythrocyte lysis buffer (155 mM NH_4_Cl, 10 mM NaHCO_3_ and 100 mM EDTA) at 1:14 ratio for 10 minutes with gentle inversion to remove erythrocytes. Then, cells were washed twice with PBS/azide (SantaCruz, Cat. sc-296028) and transferred to U-Bottom 96-well polypropylene plates (Thermo, Cat. 267245) for staining with antibodies against membrane antigens.

The resuspended cells were stained with the Fixable Violet Dead Cell Stain Kit (Thermo, Cat. L34961) to exclude dead cells. To minimize nonspecific binding, True-Stain Monocyte Blocker (Biolegend, Cat. 426103) and BD Horizon Brilliant Stain Buffer Plus (BD Biosciences, Cat. 566385) were used. Cells were stained with anti-mouse antibody mCD45 (30-F11, BD Biosciences, Cat. 553080) and the following anti-human antibodies: hCD45 (2D1, Biolegend, Cat. 368538); CD19 (HIB19, Cytek Biosciences, Cat. 60-0199-t100); CD56 (NCAM16.2, BD Biosciences, Cat. 564058); CD16 (3G8, Biolegend, Cat. 302042); CD3 (SK7, Biolegend, Cat. 344828); CD4 (SK3, Cytek Biosciences, Cat. R7-20041); CD8 (SK1, BD Biosciences, Cat. 612889); CD45RA (5H9, BD Biosciences, Cat. 740315); CD25 (BC96, Cytek Biosciences, Cat. R7-20585); HLA-DR (L243, BD Biosciences, Cat. 753688); CD14 (61D3, Cytek Biosciences, Cat. 65-0149-T100); CD11c (B-ly6, BD Biosciences, Cat. 569251); CD123 (6H6, Thermo, Cat. 62-1239-42); and CD169 (7-239, Biolegend, Cat. 346004).

After staining, cells were washed and fixed with 2% paraformaldehyde solution in PBS (SantaCruz, Cat. sc-281692) for 10 minutes at room temperature. After a final wash, cells were resuspended in PBS and stored overnight at 4°C prior to acquisition. Flow cytometric data were acquired on an Aurora spectral flow cytometer (Cytek Biosciences) configured with five lasers (355, 405, 488, 561 and 640 nm) and analyzed with FlowJo software version 10 (BD Life Sciences) according to a gating strategy that excluded murine CD45^+^ cells, identified human CD45^+^ leukocytes, and subsequently resolved major lymphoid and myeloid lineages (**Figure S1)**. This approach enabled consistent identification of T cells (CD3^+^), B cells (CD19^+^), NK cells (CD3^−^, CD19^−^ and CD56^+^ and/or CD16^+^), and a combined DC/monocyte population composed of pDCs (CD3^−^, CD19^−^, CD123^+^, CD45RA^+^), monocytes (CD3^−^, CD19^−^, CD4^mid^), and cDCs (CD3^−^, CD19^−^, CD56^−^, CD16^−^, CD11c^+^, HLA-DR^+^, CD123^−^, CD4^−^).

### 2.4. Infection of Hu-FLT3 and Hu-NSG mice

To generate X4-tropic HIV-1_NL4-3_ viral stock, 1E7 HEK-293T/17 cells (ATCC, Cat. CRL-11268) cultured in RPMI-1640 medium (Thermo, Cat. 21875-034) supplemented with 10% FBS (Thermo, Cat. 10270-106) and 1% Penicillin/Streptomycin (Thermo, Cat. 15070063) were transfected with 15 µg of pNL4-3 plasmid (US National Institute of Health (NIH) AIDS Research and Reference Reagent Program, Cat. 114) using X-tremeGENE 9 DNA Transfection Reagent (Merck, Cat. XTG9-RO) and following manufacturer’s instructions. Supernatants with viral particles were collected at 48h post-transfection, filtered (Millex HV, 0.45μm; Millipore, Cat. SLHV013SL), concentrated in PBS with LentiX reagent (Takara, Cat. 631232) and stored at −80°C. Viral stock was titrated by limiting serial dilution on the TZM-bl cell line that contains an HIV-1 long terminal repeat linked to a luciferase reporter gene (NIH AIDS Research and Reference Reagent Program, Cat. ARP5011). Luciferase expression measured with Nano-Glo Luciferase Assay System (Promega, Cat. N1110) was used to calculate TCID_50_ by the Spearman-Karber method.

Mice were either intravenously infected with 17,500 TCID_50_ of HIV_NL4-3_ virus (n = 36) or left uninfected (control group intravenously inoculated with PBS, n = 8). Body weight, HIV-1 plasma viral load and blood immunophenotype were monitored weekly and tissues were collected at final time point.

### 2.5. Administration of antiretroviral therapy to mice

As detailed above, at 5 weeks post-infection, a sub-group of the HIV-infected mice (n = 12) were placed on an ART regime consisting of a triple-combination of raltegravir (RAL, 0.6mg/g), emtricitabine (FTC, 1.5mg/g) and tenofovir disoproxil fumarate (TDF, 1.56mg/g) (Fisher Scientific, Cat. 15731799, Cat. 15791789 and Cat. 15688725, respectively). ART was absorbed in rodent chow by in house protocol (Inotiv, 2194) where a single layer of food was placed inside a sterile plastic bag with the drugs at the final concentration in 50% ethanol. The closed bag was inversed gently to achieve full absorption of the liquid phase. Then the ART-containing chow was air dry for 2 days inside the biosafety cabinet. Food was generated each week considering that each animal eats 3 gr food per day and changed two times a week. At 11 weeks post-infection, ART was discontinued, and mice were returned to standard, ART-free pelleted feed.

### 2.6. Tissue immunohistochemical analysis

Samples of spleen, sternum, kidney, liver, lung and -when found-thymus were collected and immediately fixed in neutral-buffered formalin (10%) and embedded in paraffin within 72h of fixation, following standard protocols. Sections were obtained and stained with Hematoxylin-Eosin (H/E) and evaluated by a board-certified pathologist. Additional sections were processed for immunohistochemistry (IHC) with an automatic immunostainer (Bond RX, Leica Biosystems). Briefly, slides underwent heat-induced epitope retrieval at pH 9, incubated with anti-HLA (Abcam, Cat. ab52922) or human CD11c (Abcam, Cat. ab52632) antibodies, and signal was detected through the Novolink Polymer Detection System (RE7280-CE, Leica Biosystems). Human specificity of the markers was confirmed by ruling out cross-reaction with murine antigens using non-humanized mouse tissues, which consistently yield negative results. Therefore, in the context of human-engrafted mice, any HLA positive cell at the tissue level represents either engrafted cells within the bone marrow, or lymphoid/myeloid cells that entered the bloodstream and engrafted other tissues. IHC slides were digitalized using Motic EasyScan One slide scanner at 40x magnification.

Detection of HIV-RNA within tissues was performed through RNAscope *in situ* hybridization (ISH) (Biotechne, Cat. 322150) using Bond RX immunostainer (Leica Biosystems) following manufacturer instructions, using probe 425538 targeting HIV1-CladeB genome, and 2.5 LS red assay reagents. ISH slides were digitalized at 80x magnification.

Quantification of positive cells in the tissues for corresponding markers was performed using QuPath software and expressed as % of positive cells within the tissues (24).

### 2.7. Measurement of HIV-1 viral load in blood and proviral DNA in blood and tissues

We used 30µl of plasma for HIV-RNA quantification using RealTime HIV-1 assay (Abbott Laboratories, 2G3110) with Abbott m2000 RealTime System according to the standard protocol.

Bone marrow (from femur) and spleens collected in FBS (Thermo, 10270-106) supplemented with 10 % Dimethyl Sulfoxide (DMSO, Merck, D2650) were frozen until mechanical disaggregation with cell strainers with 40 µm pore size (VWR Cat. 734-0002). Blood cells and spleen cells were treated with erythrocyte lysis buffer to reduce background as explained for immunophenotyping. After washing with cold PBS, dried cell pellets from blood, spleen or bone marrow were lysed in lysis buffer (10mM Tris-HCl, pH=9, 0.1 % Triton-X100 and 400 µg/ml Proteinase K) at 56°C during 16h for DNA extraction at concentration of 50,000cells/µl in a minimum amount of 20µl. Total HIV-DNA was measured in duplicate to quantify the frequency of proviral DNA based on primer/probe set annealing to the HIV-GAG region. The cellular RPP30 gene was measured in parallel to normalize sample input to the number of cells. Quantifications were performed by droplet digital polymerase chain reaction (ddPCR) using a QX100 Droplet Digital PCR system (Bio-Rad) as previously described (25). All probes were double-quenched 6-carboxyfluorescein (FAM)-ZEN-Iowa Black FQ (Integrated DNA Technologies). Generated QLP files were analyzed with QuantaSoft software version 1.6.6.0320 (Bio-Rad). Replicate wells were merged for quantification, and thresholds were set manually based on amplitude of negative and positive populations. In those samples lacking detection of positive droplets, LOQ was calculated as the expected result if one single droplet was positive, and LOQ/2 was used as input value for analysis.

### 2.8. Statistical analysis

Continuous variables are presented as box-and-whisker plots, and differences between groups were assessed using the Mann–Whitney U test. Longitudinal measurements of peripheral blood humanization (percentage and absolute numbers of human CD45^+^ cells) and immune cell subset composition were summarized descriptively across time points. Locally estimated scatterplot smoothing (LOESS) curves were used to visualize longitudinal trends. Differences between strains were subsequently analyzed calculating the Area Under the Curve (AUC) of the plasma RNA dynamics and using a two-way ANOVA, with treatment group and mouse strain as factors. For immunohistochemistry (human HLA and human CD11c) beta regression models were used to evaluate differences between groups and strains.

All analyses were performed using GraphPad Prism (GraphPad Software) and R (R Foundation for Statistical Computing). A two-sided significance threshold of P < 0.05 was used for inferential tests when performed.

## 3. Results

### 3.1. Myeloid and NK cell populations were sustained long-term in humanized FLT3 mice

In order to evaluate humanized FLT3 mice as experimental model for HIV-1 infection and treatment assessment (**Figure 1A**), we first characterized peripheral human immune cell reconstitution in Hu-FLT3 and Hu-NSG mice prior to HIV-1 infection in all groups (**Figure 1B**). We performed multiparametric spectral flow cytometry to identify major lymphoid and myeloid subsets within the human CD45^+^ compartment, including T cells, B cells, NK cells, and a combined DC/monocyte population composed of pDCs, monocytes and cDCs (**Figure S1**).

**Figure 1.**
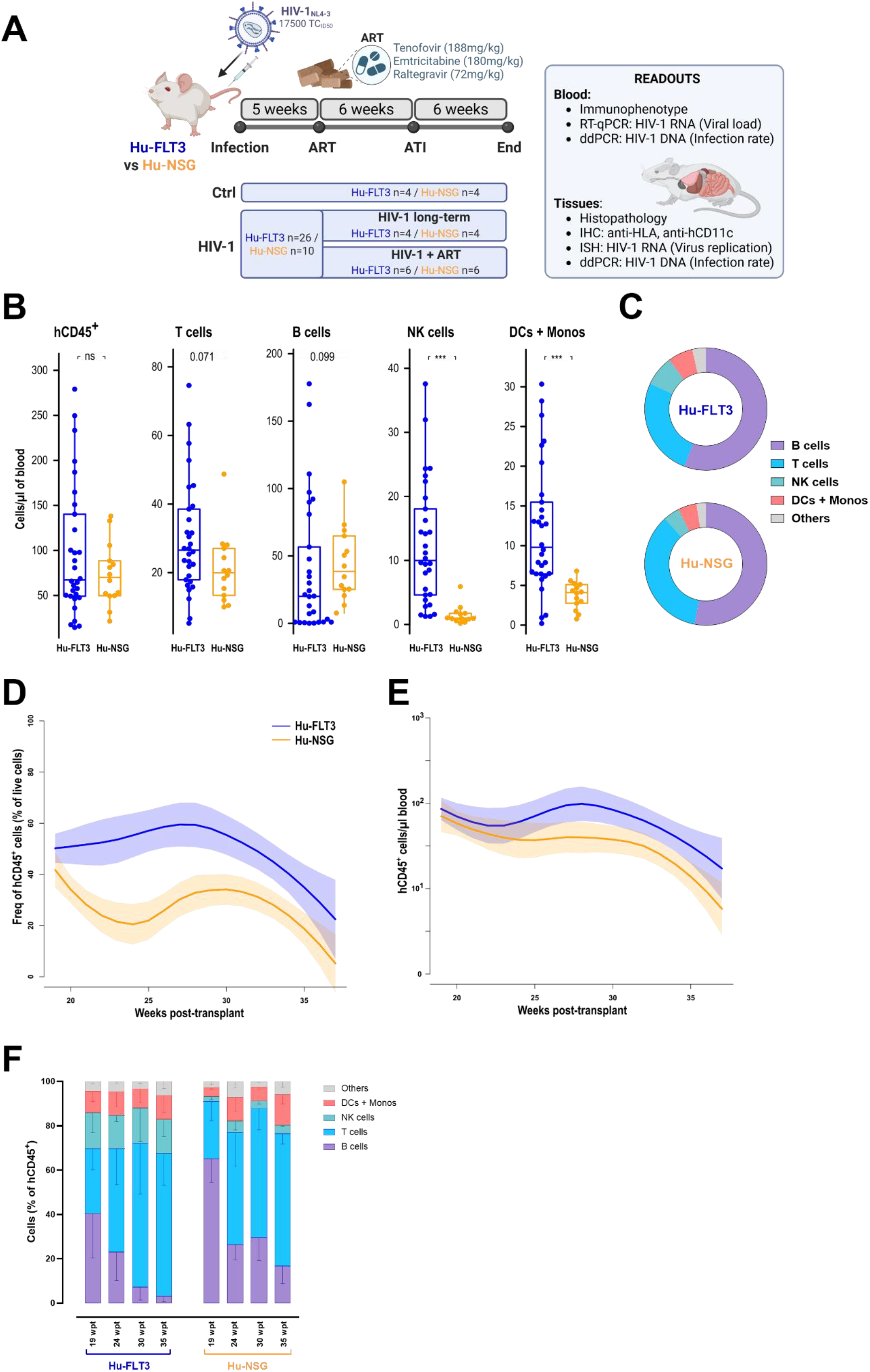
Human immune cell reconstitution in peripheral blood and tissues of Hu-FLT3 and Hu-NSG mice. **A**. Schematic representation of the experimental design. NSG and FLT3 mice were transplanted with human cord blood–derived CD34^+^ cord-blood cells from multiple independent donors and allocated into three experimental groups: Uninfected mice (**Ctrl**) received PBS and were followed until the study endpoint (Hu-FLT3 n = 4; Hu-NSG n = 4); **HIV-1** mice were intravenously infected with HIV-1_NL4-3_ and monitored for 5 weeks (short-term infection; Hu-FLT3 n = 26; Hu-NSG n = 10); a subgroup of these mice were monitored for 12 additional weeks (long-term infection; Hu-FLT3 n = 4; Hu-NSG n = 4); and a subgroup of infected mice (**HIV-1+ART**) initiated oral ART for 6 weeks, followed by analytical treatment interruption (ATI) for 6 weeks to assess viral rebound (Hu-FLT3 n = 6; Hu-NSG n = 6). **B**. Absolute numbers of human CD45^+^ cells, T cells, B cells, NK cells, and combined DCs/monocytes in peripheral blood at the earliest time point prior to infection, between week 15 and week 19 post-transplant. Each data point represents an individual animal, displayed as a box-and-whisker plot (median and interquartile range). Hu-FLT3: n = 30 (from five donors, in blue). Hu-NSG: n = 14 (from two donors, in orange). Statistical comparisons were performed using Mann-Whitney U test and significance is indicated as non-significant (ns), or P < 0.001 (***). **C**. Frequencies of T cells, B cells, NK cells, and combined DCs/monocytes in peripheral blood from samples in panel B. **D**. Longitudinal frequencies of human CD45^+^ cells in peripheral blood of Hu-FLT3 and Hu-NSG mice in control group of uninfected mice from week 19 to week 36 post-transplant. Engraftment levels are shown as percentage of total leukocytes (human CD45^+^ cells), with LOESS-smoothed trends and 95 % confidence bands displayed. Hu-FLT3: n = 4 (from two donors). Hu-NSG: n = 4 (from two donors). **E**. Absolute numbers of human CD45^+^ cells in peripheral blood of Hu-FLT3 and Hu-NSG mice from control group along week 19 to week 36 post-transplant. Graph represents LOESS-smoothed curves with 95% confidence bands. **F**. Longitudinal proportions of major human immune cell subsets in peripheral blood of mice in control group, expressed as percentage of hCD45^+^ cells at 19, 24, 30, and 35 weeks post-transplant. Subsets include B cells (purple, CD19^+^), T cells (blue, CD3^+^), NK cells (green, CD3^−^ and CD56^+^ and/or CD16^+^), and a combined DC/monocyte population (red) comprising monocytes (CD4^mid^), conventional dendritic cells (CD11c^+^ HLA-DR^+^ CD123^−^ CD4^−^), and plasmacytoid dendritic cells (CD123^+^ CD45RA^+^).

At the earliest experimental time point before infection (15–19 weeks post-transplant), Hu-FLT3 mice exhibited significantly higher absolute numbers of NK cells and combined DC/monocyte populations compared with Hu-NSG mice, whereas absolute numbers of T cells and B cells were comparable between strains (**Figure 1B**). Analysis of relative cell frequencies showed that NK cells and DC/monocyte populations represented a substantially larger fraction of the human CD45^+^ compartment in Hu-FLT3 mice than in Hu-NSG mice (six-fold and three-fold smaller, respectively) (**Figure 1C**). We next assessed the stability of human immune cell engraftment over time in uninfected control mice. From week 19 to week 36 post-transplant, the proportion of human CD45^+^ cells among total peripheral blood live cells remained stable in both strains with a progressive decay from week 30 post-transplant (**Figure 1D**). Throughout this period, Hu-FLT3 mice consistently maintained higher mean levels of circulating human CD45^+^ cells than Hu-NSG mice, as reflected by both relative frequencies and absolute cell counts (**Figure 1D-E**). Humanization outcomes varied across individual donors and recipient animals in both strains, highlighting donor-dependent and inter-individual heterogeneity in immune reconstitution (**Figure S2A**). Despite this variability, Hu-FLT3 mice reproducibly displayed higher frequencies of NK cells and combined DC/monocyte populations at all analyzed time points, whereas these compartments remained markedly reduced in Hu-NSG mice (**Figures 1F, S2B**). These results demonstrate that enhanced innate immune cell reconstitution in Hu-FLT3 mice was robust across multiple donors, despite biologically relevant variation between humanization sources.

To determine whether enhanced peripheral engraftment of human cells extended to tissues, we examined lymphoid and non-lymphoid organs by immunohistochemistry (IHC) in the non-infected control mice at the end of the study (36 weeks post-transplant) (**Figure 2A**). Human HLA^+^ cells were abundantly detected in bone marrow and spleen sections in both strains and accounted for the majority of hematopoietic cells within bone marrow (80 to 100% of total hematopoietic cells). Human CD11c^+^ cells, used as a surrogate marker for myeloid populations, were also detected in these tissues at lower frequency.

**Figure 2.**
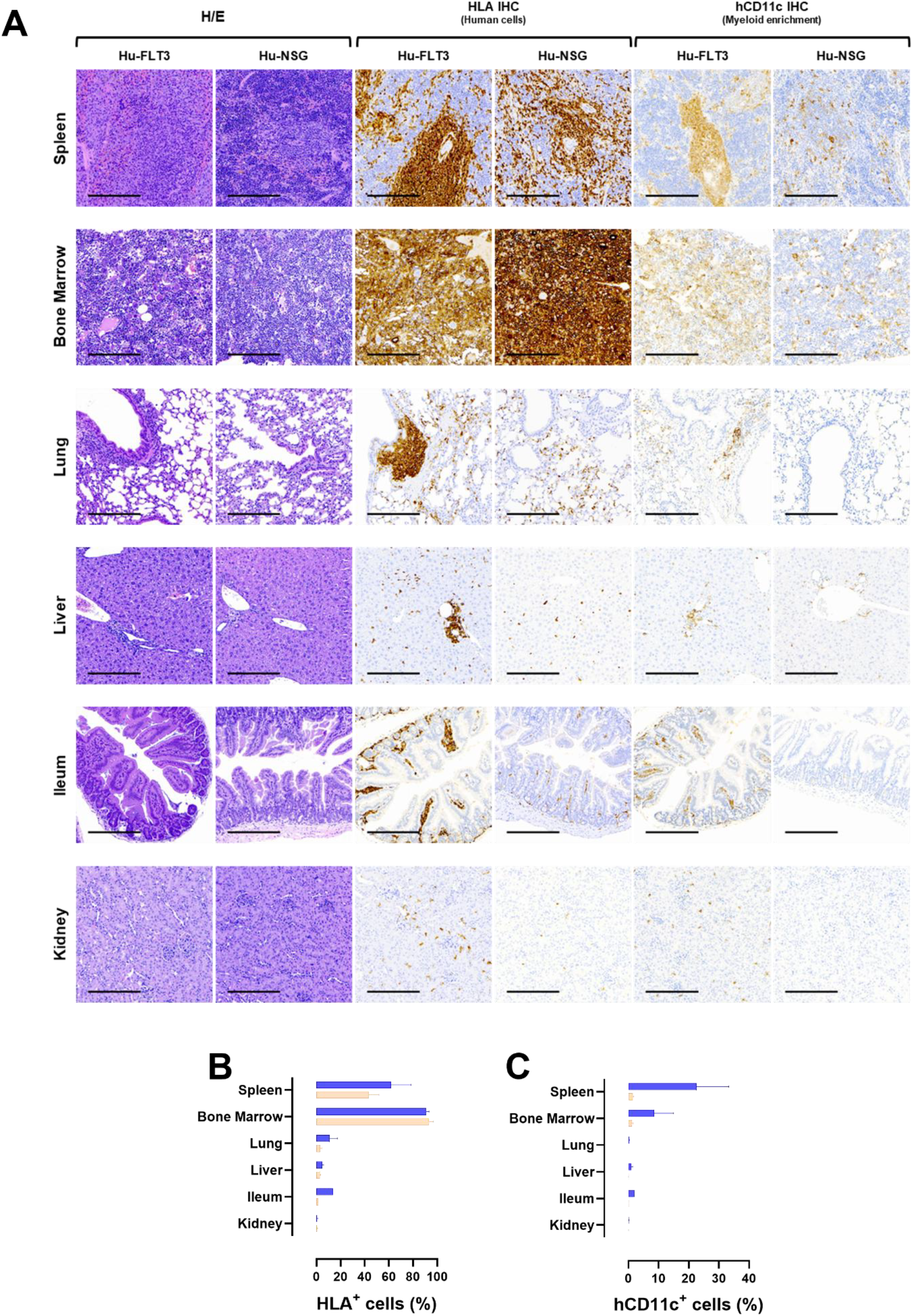
Human immune cell engraftment in non-lymphoid tissues of Hu-FLT3 and Hu-NSG mice from control group. Tissue sections from spleen, bone marrow lung, liver, ileum, and kidney were stained with hematoxylin and eosin (H/E), anti-HLA to identify human hematopoietic cells, and anti-human CD11c to detect myeloid cell subsets at week 36 post-transplant. **A**. Representative histological images of indicated tissues. Scale bar = 200 µm. **B**. Quantification of human HLA-positive and human CD11c-positive cells. Data represents the percentage of positive cells within the analyzed tissue, as determined using QuPath image analysis software.

Remarkably, we also detected human immune cells in non-lymphoid organs, including liver, lung, ileum, and kidney, in expected histological locations such as portal tracts in the liver. In the lung, organized bronchus-associated lymphoid tissue–like (BALT-like) follicles populated by human immune cells were observed exclusively in Hu-FLT3 mice (**Figure 2A**), indicating enhanced tissue-level immune organization in this mouse model. Quantitative image analysis confirmed that tendency to find higher levels of human HLA^+^ and CD11c^+^ signal in Hu-FLT3 tissues compared with Hu-NSG tissues (**Figures 2B, C**).

Together, these results showed that Hu-FLT3 mice supported sustained and enhanced reconstitution of human innate immune cells in both blood and tissues over prolonged periods post-transplant without compromising lymphocytic population frequencies. Although humanization outcomes varied across donors and individuals, the expanded NK and myeloid compartments observed in Hu-FLT3 mice were consistent across multiple donors, underscoring the value of multi-donor humanization strategies for generating robust and biologically representative *in vivo* models.

### 3.2. Hu-FLT3 and Hu-NSG mice supported productive HIV-1 infection in blood and tissues

We and others have previously shown that humanized NSG mice are susceptible to infection with HIV-1 (17). Here, we next evaluated whether Hu-FLT3 mice also supported productive HIV-1 infection and compared early infection dynamics with the Hu-NSG model (**Figure 1A**, HIV group). Both strains were intravenously inoculated with 17,500 TCID_50_ of HIV-1_NL4-3_ and monitored longitudinally for virological and immunological parameters.

At 1 week post-infection (wpi), both Hu-FLT3 and Hu-NSG mice developed detectable plasma viremia (**Figure 3A**). Plasma viral loads increased during the first five weeks of infection in both strains, with Hu-FLT3 mice exhibiting higher mean levels of viremia at 5 wpi (1.9 × 10^5^ copies/ml) compared with Hu-NSG mice (5.8 × 10^4^ copies/ml).

**Figure 3.**
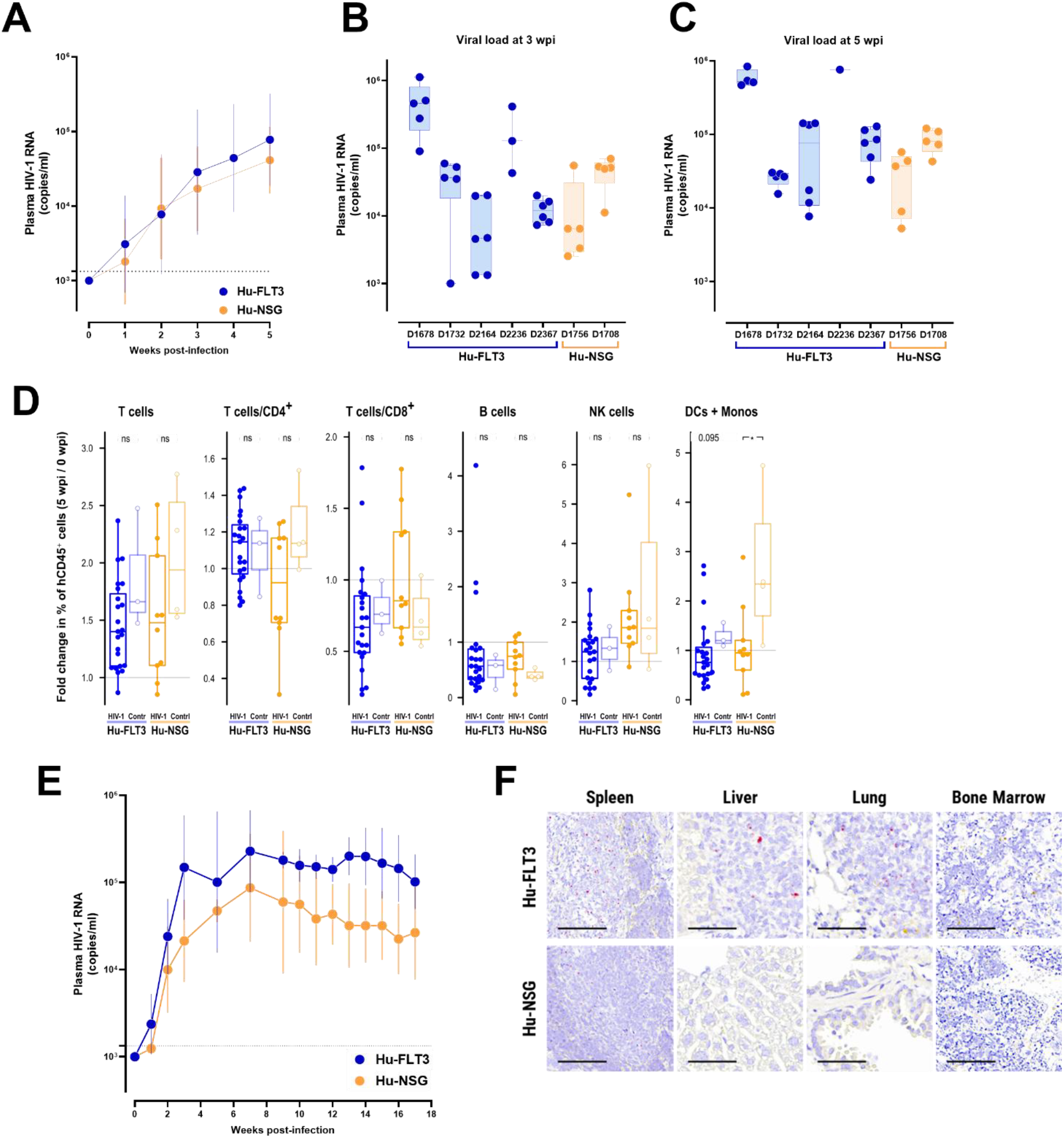
Hu-FLT3 and Hu-NSG humanized mice support productive HIV-1 infection. Blood samples were collected weekly from Hu-FLT3 (blue, n = 26) and Hu-NSG (orange, n = 10) infected mice. **A**. Plasma HIV-1 RNA levels in HIV-1 mice from week 0 to week 5 post-infection. Viral loads are displayed as geometric mean ± geometric SD. The lower limit of detection (LOD) is 1.3 × 10^3^ copies/ml is indicated with a dotted horizontal line; undetectable samples were assigned a value of 10^3^ copies/ml. **B**. Plasma HIV-1 RNA levels at 3 wpi stratified by donor in Hu-FLT3 (blue, five donors) and Hu-NSG (orange, two donors) mice. Viral loads are shown as individual values with box-and-whisker plots. **C**. Plasma HIV-1 RNA levels at 5 wpi stratified by donor in Hu-FLT3 (blue, five donors) and Hu-NSG (orange, two donors) mice. Viral loads are shown as individual values with box-and-whisker plots. **D**. Changes in major human immune cell subsets in peripheral blood upon infection. Fold change values represent the ratio of the frequency (% of hCD45^+^ cells) at 5 wpi relative to baseline (0 wpi) for T cells (including CD4^+^ and CD8^+^ T-cell subsets), B cells, NK cells, and the combined DC/monocyte populations. Groups compared include Hu-FLT3 HIV-1 (n = 21, dark blue), Hu-FLT3 Ctrl (n = 3, non-infected control group, light blue), Hu-NSG HIV-1 (n = 10, dark orange), and Hu-NSG Ctrl (n = 4, non-infected control group, light orange). Statistical comparisons were performed using Mann-Whitney U test; significance is indicated as non-significant (ns) or P < 0.05 (*). **E**. Plasma HIV-1 RNA levels in subgroup of infected mice from week 0 to week 17 post-infection. Viral loads are displayed as geometric mean ± geometric SD comparing Hu-FLT3 (blue, two donors, n = 4) and Hu-NSG (orange, two donors, n =4) infected mice. The lower limit of detection (LOD) is 1.3 × 10^3^ copies/ml, indicated with a dotted horizontal line; undetectable samples were assigned a value of 10^3^ copies/ml. **F**. Detection of HIV-1 RNA in tissues of infected Hu-FLT3 and Hu-NSG mice at week 17 post-infection. Representative RNAscope *in situ* hybridization images from spleen, liver, lung, and bone marrow. HIV-1 RNA signal is shown in pink. Scale bar = 50µm.

To assess the influence of donor-to-donor variability on infection outcomes, we stratified plasma viral loads by donor at 3 and 5 wpi and observed marked variability in viral loads among donors (**Figure 3B, C**). Despite this heterogeneity, mice engrafted with cells from all donors supported productive infection, and no animals from either strain exhibited low or undetectable plasma viremia at these early time points. We did not observe a correlation between plasma viral load levels and any immunophenotypic profile, emphasizing the complex contribution of donor-dependent human immune reconstitution to infection dynamics.

We then assessed early changes in major circulating human immune cell populations following HIV-1 infection. We calculated fold changes in T cells (including CD4^+^ and CD8^+^ subsets), B cells, NK cells, and combined DC/monocyte populations relative to baseline at 5 wpi (**Figures 3D**). Overall, patterns of immune cell variation following infection were similar between Hu-FLT3 and Hu-NSG mice. T-cell frequencies increased less in infected mice than in uninfected controls, consistent with early effects of HIV-1 infection on the human T-cell compartment. Detailed analysis of CD4^+^ and CD8^+^ T-cell subsets showed modest changes during early infection (**Figures 3D**). CD8^+^ T-cell frequencies decreased relative to baseline across infected groups, whereas CD4^+^ T-cell frequencies showed small increases in both strains. The proportion of DC/monocyte subpopulations was continuous upon infection while it increased in uninfected controls, as also observed in the longitudinal analysis of peripheral immune cell dynamics (**Figure S3**). In contrast, B-cell and NK-cell frequencies remained stable during the first five weeks of infection in both strains (**Figures 3D**). While T-cell frequencies gradually increased and B-cell frequencies declined similarly in both strains, NK cells and DC/monocyte populations were consistently present at higher levels in Hu-FLT3 mice than in Hu-NSG mice at all evaluated time points (**Figure S3**).

Both humanized strains supported productive HIV-1 infection across the 17-week follow-up period, with sustained plasma viremia observed in untreated animals and no evidence of spontaneous viral clearance in either strain (**Figure 3E**). To determine whether productive viral replication extended beyond the bloodstream, we analyzed tissue-associated viral RNA by RNAscope *in situ* hybridization. At 17 wpi, we detected HIV-1 RNA-positive cells predominantly in spleen, with lower-level signal observed in other tissues where human immune cells were present, including liver portal spaces and bronchus-associated lymphoid tissue–like structures in the lung (**Figure 3F**). The intensity and distribution of viral RNA signal were greater in Hu-FLT3 mice than in Hu-NSG mice, in which tissue-associated viral RNA was sparse or undetectable with this technology. We additionally detected HIV-1 RNA in tissues of Hu-FLT3 mice at 5 wpi, confirming early dissemination of infection to peripheral organs (**Figure S4**).

Together, these findings demonstrate that both Hu-FLT3 and Hu-NSG mice were permissive to HIV-1 infection and supported sustained viral replication in blood and tissues during early infection. Hu-FLT3 mice exhibited higher plasma viremia and more abundant tissue-associated viral RNA, consistent with their expanded and durable human innate immune cell repertoire and emphasizing the relevance of this model for studying early HIV-1 dissemination across multiple donor backgrounds.

### 3.3. Hu-FLT3 and Hu-NSG mice suppressed viremia under ART and exhibited viral rebound upon treatment interruption

We next evaluated the response of Hu-FLT3 and Hu-NSG mice to antiretroviral therapy (ART) and their capacity to sustain HIV-1 persistence after treatment interruption. A subgroup of HIV-1–infected mice from both strains initiated oral ART at 5 wpi and was monitored through 17 wpi, in parallel with a subgroup of untreated infected animals (**Figure 1A**). Initiation of ART induced a rapid decline in plasma viremia in Hu-FLT3 and Hu-NSG mice, and all treated animals reached undetectable HIV-1 RNA levels within four weeks of therapy (**Figure 4A**). ART was well tolerated in both models, and no spontaneous viral suppression was observed in untreated controls.

**Figure 4.**
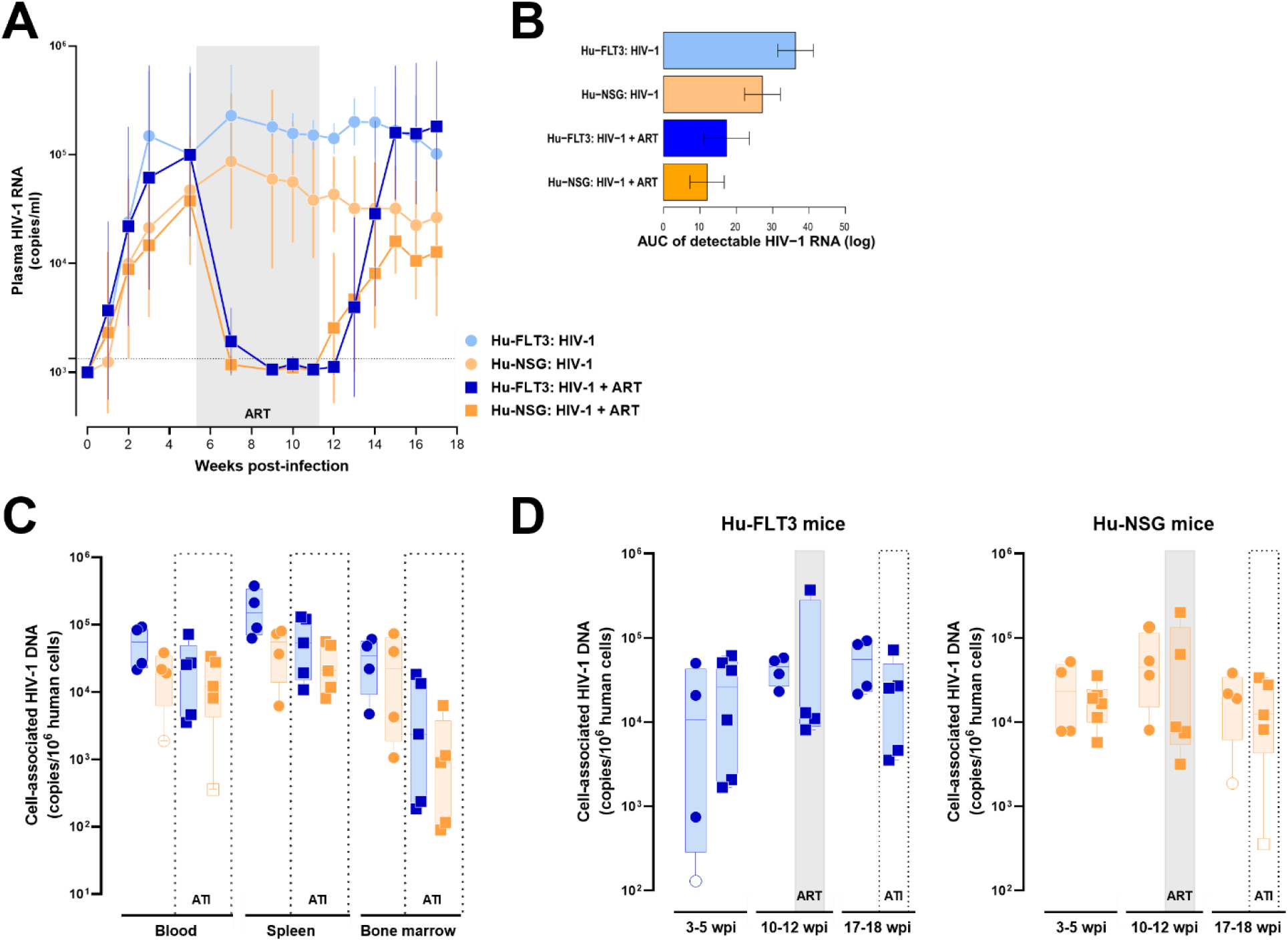
Hu-FLT3 and Hu-NSG mice suppress plasma HIV-1 viremia under oral antiretroviral treatment (ART) and exhibit uniform viral rebound following treatment interruption (ATI). Hu-FLT3 (blue) and Hu-NSG (orange) mice were intravenously infected with 17,500 TCID_50_ HIV-1_NL4-3_. After 5 weeks, mice were placed under suppressive ART (squares; n = 6 per strain) via food for 6 weeks (5–11 wpi) in comparison to control group (circles; n = 4 per strain) that stayed with normal food. All animals were monitored weekly until 17 wpi. **A**. Longitudinal plasma HIV-1 RNA kinetics measured by qRT-PCR. Viral loads are displayed as geometric mean ± geometric SD. ART duration is highlighted by a grey shaded region. The lower limit of detection (LOD) is 1.3 × 10^3^ copies/ml and indicated with a dotted horizontal line; undetectable samples were assigned a value of 10^3^ copies/ml. **B**. Viral load in plasma has been calculated as the area under the curve (AUC) for each mouse over 0– 17 wpi and differences were analyzed using a two-way ANOVA. Values are displayed as mean AUC with CI 95 %. **C**. Cell associated HIV-1 DNA measured by ddPCR in blood, spleen, and bone marrow at 17 wpi, to compare Hu-FLT3 mice on ART (blue square, n = 5), Hu-FLT3 without treatment (blue circle, n = 4), Hu-NSG on ART (orange square, n = 5), and Hu-NSG without treatment (orange circle, n = 4). Data are shown as individual values and open symbols indicate values below the detectable threshold. **D**. HIV-1 DNA quantified by ddPCR in whole-blood cells at early infection before starting ART (3-5 wpi), before ART interruption (10-12 wpi) and after viral rebound at end point (17-18 wpi). Data are shown as individual values and open symbols indicate values below the detectable threshold.

Following analytical treatment interruption (ATI) at 11 wpi, plasma viremia became detectable again in all ART-treated mice from both strains within four weeks post-ATI (**Figure 4A**). Although viral rebound occurred uniformly across treated animals, Hu-FLT3 mice reached higher rebound plasma HIV-1 RNA levels compared with Hu-NSG mice (**Figures 4A**), approaching those observed during early infection. The analysis of the area under the plasma RNA dynamics (AUC) using a two-way ANOVA reflected the expected difference between ART group and non-treated animals (pval = 2.25 × 10^−5^) (**Figures 4B**). It also showed a significant difference between the two strains (pval = 0.022) and we concluded that this strain effect is consistent in treated and non-treated groups because the interaction term is not significant (pval = 0.5).

To further assess viral persistence across compartments, we quantified cell-associated HIV-1 DNA in blood, spleen, and bone marrow at end point (17 wpi, with detectable viremia) using droplet digital PCR. HIV-1 DNA was detectable in all analyzed tissues in both strains, and comparable to untreated animals (**Figure 4C**). While variability was observed between individual animals, Hu-FLT3 mice tended to exhibit higher levels of tissue-associated HIV-1 DNA than Hu-NSG mice, although these differences were not subjected to statistical inference. We then examined the dynamics of blood-associated HIV-1 DNA over time and measured total HIV-1 DNA levels during early infection (3-5 wpi), during suppressive ART (10-12 wpi), and at the final study endpoint in both mouse models (**Figure 4D**). HIV-1 DNA levels remained detectable during ART and after treatment interruption, consistent with the persistence of proviral material under suppressive therapy in both strains.

To contextualize viral persistence at the tissue level, we evaluated spleen sections collected at the terminal time point in all groups of animals by immunohistochemistry and RNAscope *in situ* hybridization (**Figure 5**). HLA^+^ cells were readily detected in spleens from all infected animals in both strains, indicating sustained human hematopoietic engraftment at the end of the study (**Figures 5A-B**). Analysis of human HLA^+^ and CD11c^+^ cell frequency was performed using beta regression models including treatment and strain as predictors. We observed a significant expression increase for both HLA^+^ and human CD11c^+^ markers in the Hu-FLT3 strain compared to Hu-NSG strain. Tissue-associated HIV-1 RNA was detected in spleens of infected mice, with stronger signal in Hu-FLT3 animals compared to Hu-NSG animals, that showed almost undetectable levels, whereas uninfected controls remained uniformly negative (**Figure 5D**). Similar trends of sustained human cell engraftment were observed in bone marrow sections (**Figure S5**), although HIV-1 RNA was undetectable (data non shown).

**Figure 5.**
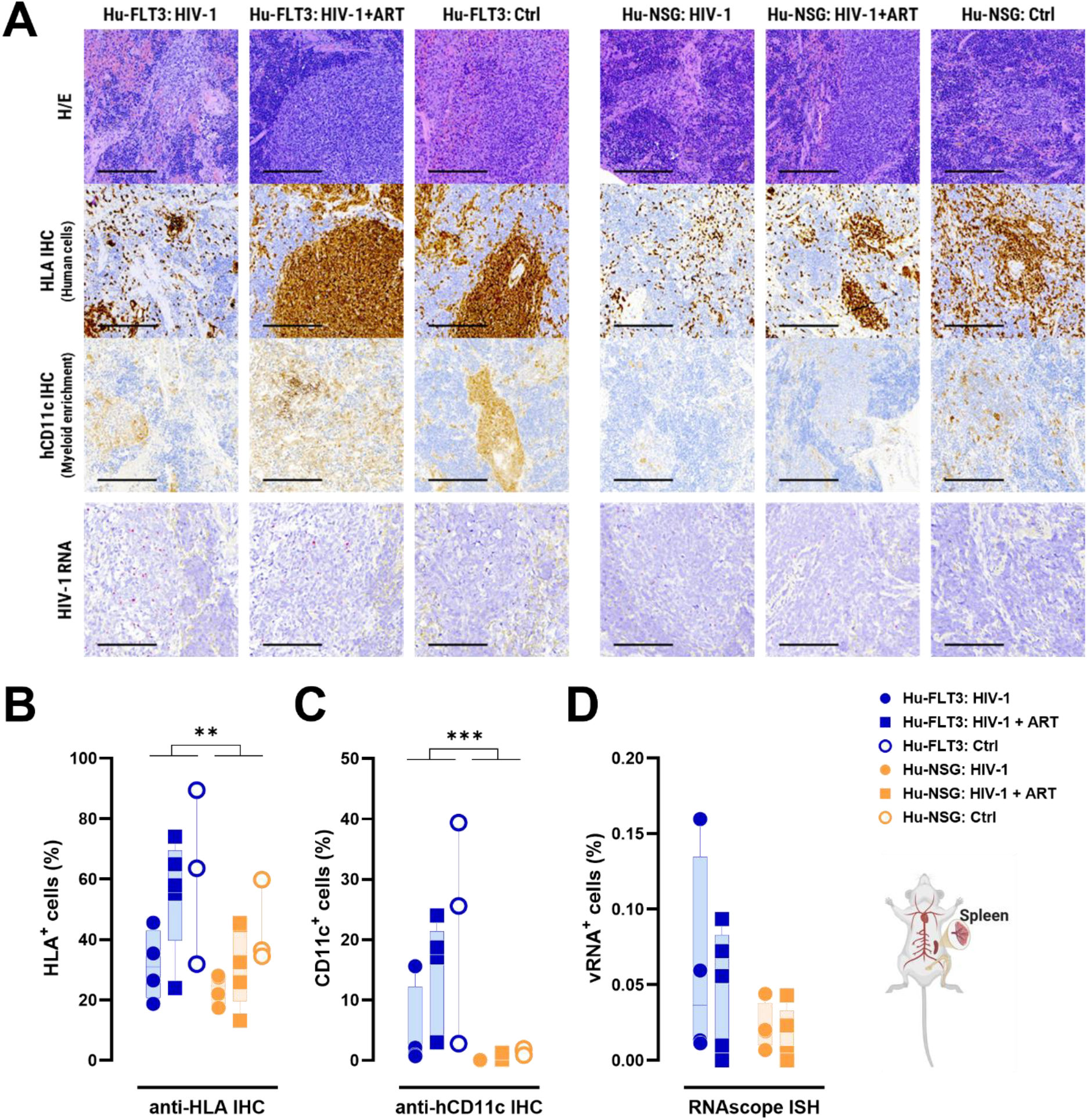
Human immune cell reconstitution and tissue-associated HIV-1 RNA in spleen of humanized mice. Hu-FLT3 and Hu-NSG mice were analyzed at the terminal timepoint (17 wpi) to compare only HIV-1 infected mice (Hu-FLT3, blue circles, n = 4; Hu-NSG, orange circles, n = 4;), HIV-1 infected on ART (Hu-FLT3, blue squares, n = 5; Hu-NSG, orange squares, n = 5), and non-infected controls (Hu-FLT3 n = 3; Hu-NSG n = 3; open circles). Spleen sections were evaluated by H/E staining, HLA and human CD11c immunohistochemistry, and RNAscope HIV-1 RNA *in situ* hybridization. Quantification was performed in QuPath software as percent positive cells for HLA, CD11c, and HIV-1 RNA within tissue. **A**. Representative histological images of spleen sections from Hu-FLT3 and Hu-NSG mice stained with indicated cellular markers (brown) and HIV-1 RNA (pink). Scale bar = 200 µm. **B**. Quantification of HLA-positive cells. **C**. Quantification of human CD11c-positive cells. **D**. Quantification of spleen-associated HIV-1 RNA by RNAscope from infected animals (uninfected controls were all negative). In **B, C** and **D**, Values for individual mice are shown and differences were evaluated using beta regression models. The significance of strain’s coefficient is indicated as P < 0.001 (***) or P < 0.01 (**).

Together, these results demonstrated that both Hu-FLT3 and Hu-NSG mice responded efficiently to oral ART, suppressed plasma viremia to undetectable levels, and uniformly exhibited viral rebound following treatment interruption. The sustained presence of proviral DNA and tissue-associated HIV-1 RNA in blood and lymphoid organs supported the establishment of persistent viral reservoirs in both models. Hu-FLT3 mice showed consistently higher rebound viremia and greater tissue-level viral and immune cell signals, highlighting the value of this model for studying HIV-1 persistence and treatment interruption in the context of an expanded human immune repertoire.

## 4. Discussion

Humanized mouse models constitute a key experimental platform for studying HIV-1 infection, persistence, and therapeutic interventions *in vivo* (26). However, widely used models present limitations regarding immune system reconstitution, cellular lineage diversity, or technical accessibility. PBMC-based models offer rapid engraftment but most are restricted by graft-versus-host disease, limiting their utility to short-term studies (27). BLT models display the most complete human immune reconstitution, including robust thymic development, but depend on fetal tissue procurement, which raises ethical, logistical, and regulatory barriers that impede broad adoption (28). On the other hand, HSC-based NSG models avoid these constraints and support long-term studies, yet they typically exhibit limited reconstitution of myeloid and NK cell compartments. The Hu-FLT3 model evaluated here addresses several of these limitations by supporting markedly improved NK and myeloid lineage development (21,22,29). We provide a comprehensive characterization of the Hu-FLT3 humanized mouse model in the context of HIV-1 infection, antiretroviral therapy, and treatment interruption, and compare its performance with the Hu-NSG model.

A central finding of this work is that Hu-FLT3 mice supported enhanced and durable reconstitution of human innate immune cell populations in both blood and tissues. Compared with Hu-NSG mice, Hu-FLT3 animals displayed expanded NK-cell and myeloid compartments that were maintained for 36 weeks post-transplant. Importantly, this enhanced immune reconstitution was observed across multiple independent donors, despite marked donor-to-donor variability in overall humanization levels and immune complexity. These results highlight both the biological relevance of multi-donor humanization strategies for generating robust and representative preclinical data. Enhanced peripheral blood immune reconstitution in Hu-FLT3 mice was paralleled by increased engraftment in the tissues. Human immune cells were widely distributed across lymphoid and non-lymphoid organs. Moreover, Hu-FLT3 mice uniquely displayed organized bronchus-associated lymphoid tissue–like structures in the lung that suggests improved immune organization at mucosal sites. This enhanced immune complexity provides an important advantage for HIV-1 studies, as both NK cells and myeloid cells contribute to sensing, restricting, or harboring HIV-1 depending on anatomical location and activation state (30,31).

Both Hu-FLT3 and Hu-NSG mice supported productive HIV-1 infection following intravenous infection, with sustained plasma viremia observed in untreated animals for up to 17 weeks. However, Hu-FLT3 animals consistently showed more abundant tissue-associated HIV-1 RNA during infection, potentially because of the expanded innate immune repertoire in these mice, a feature relevant to human tissue-resident infection. Previous studies have described that monocytes and macrophages contribute to viral replication and reservoir establishment, while DCs in tissue contribute to a broader tissue dissemination of HIV-1 (32).

Oral ART delivered via drug-containing food efficiently suppressed plasma viremia in both Hu-FLT3 and Hu-NSG mice, demonstrating that this non-invasive administration route is suitable for long-term HIV studies requiring minimal animal handling. Following analytical treatment interruption, all treated animals exhibited viral rebound, indicating the persistence of replication-competent viral reservoirs under suppressive therapy. Quantification of HIV-1 DNA in blood cells and tissues further demonstrated that both mouse models maintain proviral reservoirs in multiple anatomical sites. Hu-FLT3 mice reached higher rebound viremia than Hu-NSG mice, approaching levels observed during early infection. Tissue analyses further revealed sustained human immune cell engraftment and detectable HIV-1 RNA in spleen sections, particularly in Hu-FLT3 mice, feature particularly relevant to investigate HIV-1 persistence in tissue reservoirs (33). These results support the use of the Hu-FLT3 model for studies aimed at evaluating viral rebound dynamics that are central to treatment interruption studies in people living with HIV.

The findings of this study are subject to a series of limitations. First, donor-dependent variability in human immune reconstitution was substantial and contributed to heterogeneity in virological and immunological outcomes. While this variability reflects the natural diversity observed in human populations, it also emphasizes the need for adequate donor representation and cautious interpretation of group-level comparisons. Second, although Hu-FLT3 mice exhibit markedly improved myeloid and NK cell development, human microglia and other CNS-resident myeloid populations were not assessed, and future studies will be required to determine the suitability of the Hu-FLT3 model for investigating HIV-1 infection and persistence in the central nervous system and other immune-privileged sites (34,35). Future studies should also evaluate mucosal routes of infection to determine how broadly the Hu-FLT3 model recapitulates human HIV-1 transmission and pathogenesis (36). Finally, although tissue-associated viral RNA and proviral DNA was readily detected, functional assays of myeloid or NK cell responses were not performed and will be needed to define how these expanded lineages contribute to antiviral immunity or reservoir maintenance in Hu-FLT3 mouse model.

In summary, this work establishes the FLT3 humanized mouse model as a robust and accessible *in vivo* platform that supports enhanced human innate immune reconstitution, sustained HIV-1 replication, effective antiretroviral suppression and viral rebound upon treatment interruption. By combining improved immune lineage diversity with reproducible virological outcomes across multiple donors, together with its accessibility and absence of fetal tissue requirements, the Hu-FLT3 model provides a robust, ready-to-use tool for preclinical evaluation of HIV-1 cure interventions, including antiviral drugs, therapeutic antibodies and vaccine strategies, as wells as persistence-targeting strategies relevant to clinical development.

## Abbreviations

ART: antiretroviral therapy
cat. no.: catalogue number
cDCs: conventional dendritic cells
FLT3: NOD.Cg-*Flt3*^*em2Mvw*^ *Prkdc*^*scid*^ *Il2rg*^*tm1Wjl*^ *Tg(FLT3LG)*7Sz/SzJ mice
HIV-1: human immunodeficiency virus
Hu: humanized mouse model
HSC: hematopoietic stem cells
NSG: NOD.Cg-*Prkdc*^*scid*^ *Il2rg*^*tm1Wjl*^/SzJ mice
pDCs: plasmacytoid dendritic cells

## Acknowledgements

We are grateful to the technical contribution from Marta Margalef and Sergi Sunye from CMCiB-IGTP; Sara Monreal, Joan Puñet, Marco and A. Fernández from IGTP; Águeda Hernández from HGTIP; Mónica Pérez and Gema Garcia from IRTA-CReSA; and Lidia Garrido from IrsiCaixa. We thank prof. Matthew D Marsden (School of Medicine, University of California, Irvine, USA) for kindly sharing the protocol to prepare ART regime in rodent chow. We thank the Scientific and technical services (IrsiCaixa) for the blood sample processing and PBMCs isolation. Some figures were created with BioRender.com.

AI Use Declaration: Artificial intelligence–assisted tools (Microsoft Copilot/GPT-based language models) were used to support grammar refinement and text structuring during manuscript preparation. All scientific content, data interpretation, and final text were generated solely by the authors that take full responsibility for the accuracy and integrity of the manuscript.

## Funding sources

The authors’ laboratories were supported by funding from the Spanish Ministry of Science and Innovation (grants PID2022-139271OB-I00 and CB21/13/00063, Spain), Generalitat de Catalunya (CERCA program and 2021-SGR-00452) NIH/NIAID (1UM1AI164561-01 and 1P01AI178376-01, United States of America), and Grifols (Spain). MS was supported by the Miguel Servet Fellowship Program (CP22/00038) and I+D+I RTI-A Grant (PID2023-147320OB-I00) from the Ministry of Science and Innovation. MAB and LDS were supported by funding from NIH (R24OD036199) and by the Barbara D. Cammett Breakthrough T1D Center of Excellence in New England of the Breakthrough T1D Foundation (4-COE-2025-1751-A-N).

## Author contributions

PR-I: Conceptualization, Data curation, Formal analysis, Investigation, Methodology, Project administration, Supervision, Validation, Visualization, Writing – original draft, Writing – review and editing

PP: Data curation, Investigation, Methodology, Writing (review & editing)

FL: Investigation, Methodology, Writing (review & editing)

GC-G: Investigation, Methodology, Writing (review & editing)

AC: Data curation, Investigation, Methodology, Visualization, Writing (review & editing)

VU: Investigation, Formal analysis, Visualization, Writing – review and editing

MG-G: Investigation, Writing (review & editing)

YR-S: Investigation, Writing (review & editing)

MC: Investigation, Writing (review & editing)

JD-P: Investigation, Writing (review & editing)

LS: Methodology, Writing (review & editing)

VY: Methodology, Writing (review & editing)

MB: Methodology, Writing (review & editing)

BS: Methodology, Writing (review & editing)

MS: Conceptualization, Formal analysis, Investigation, Methodology, Validation, Writing – review and editing

JM-P: Conceptualization, Funding acquisition, Investigation, Methodology, Resources, Supervision, Validation, Writing – review and editing

## Disclosure and competing interest statements

J.M.-P. has received institutional grants and educational/consultancy fees from Gilead Sciences, Grifols, Merck Sharp & Dohme, and ViiV Healthcare, all outside the submitted work. L.S., V.Y. and B.S. are employees of The Jackson Laboratory, a not-for-profit organization commercializing the mouse strains used in these studies. Other authors declare no financial or commercial conflict of interest.

## FIGURES

**Figure S1.**
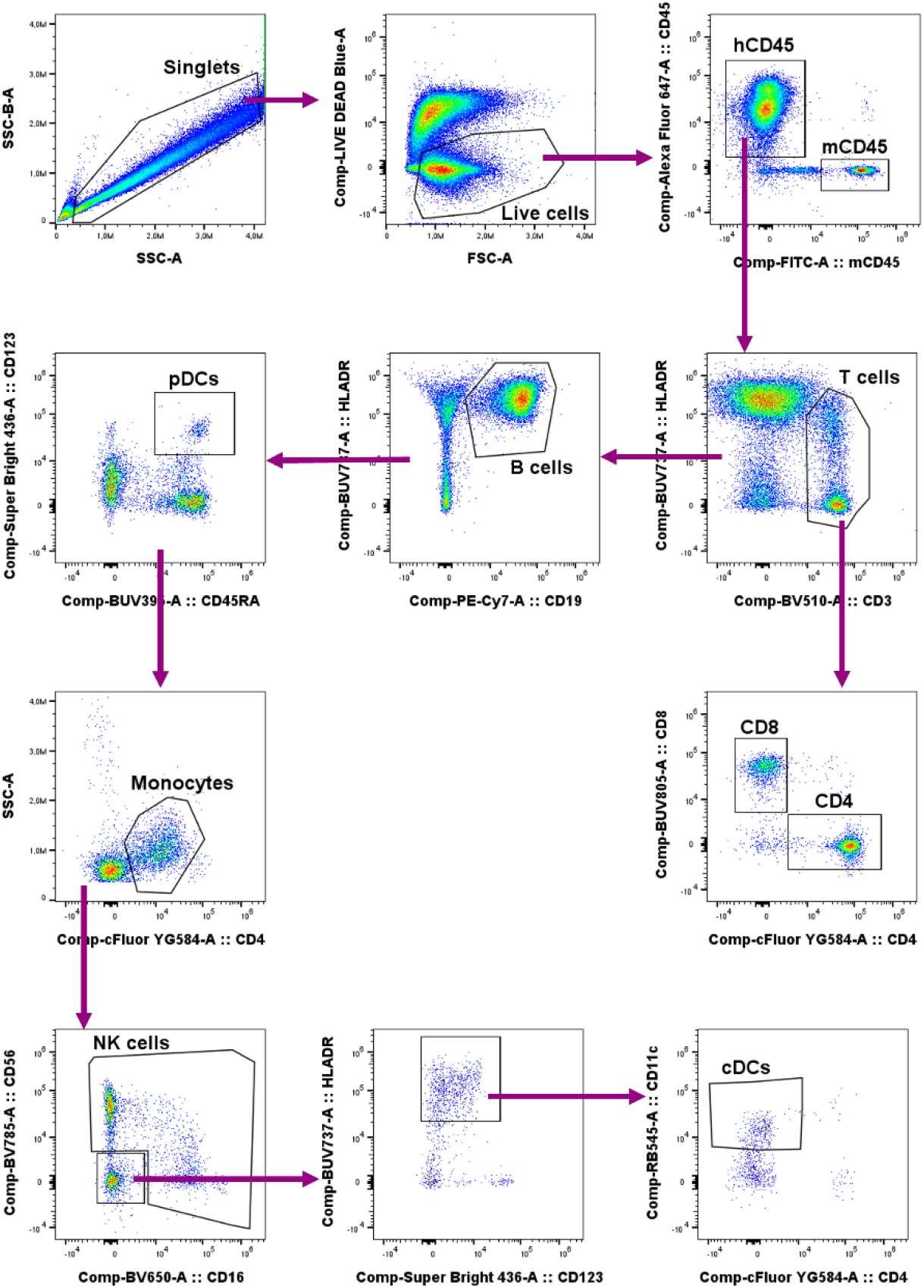
Multiparametric spectral flow cytometry gating strategy used to identify human immune cell subsets in peripheral blood of Hu-FLT3 and Hu-NSG mice. After exclusion of doublets and dead cells, murine CD45^+^ cells were removed and human CD45^+^ leukocytes were identified. Within the hCD45^+^ compartment, major lymphoid and myeloid lineages were resolved sequentially, including T cells (CD3^+^), B cells (CD19^+^), pDCs (CD123^+^, CD45RA^+^), monocytes (CD4^mid^), NK cells (CD56^+^ and/or CD16^+^), and cDCs (HLA-DR^+^, CD123^−^, CD11c^+^, CD4^−^). This gating strategy was applied consistently across all experimental timepoints.

**Figure S2.**
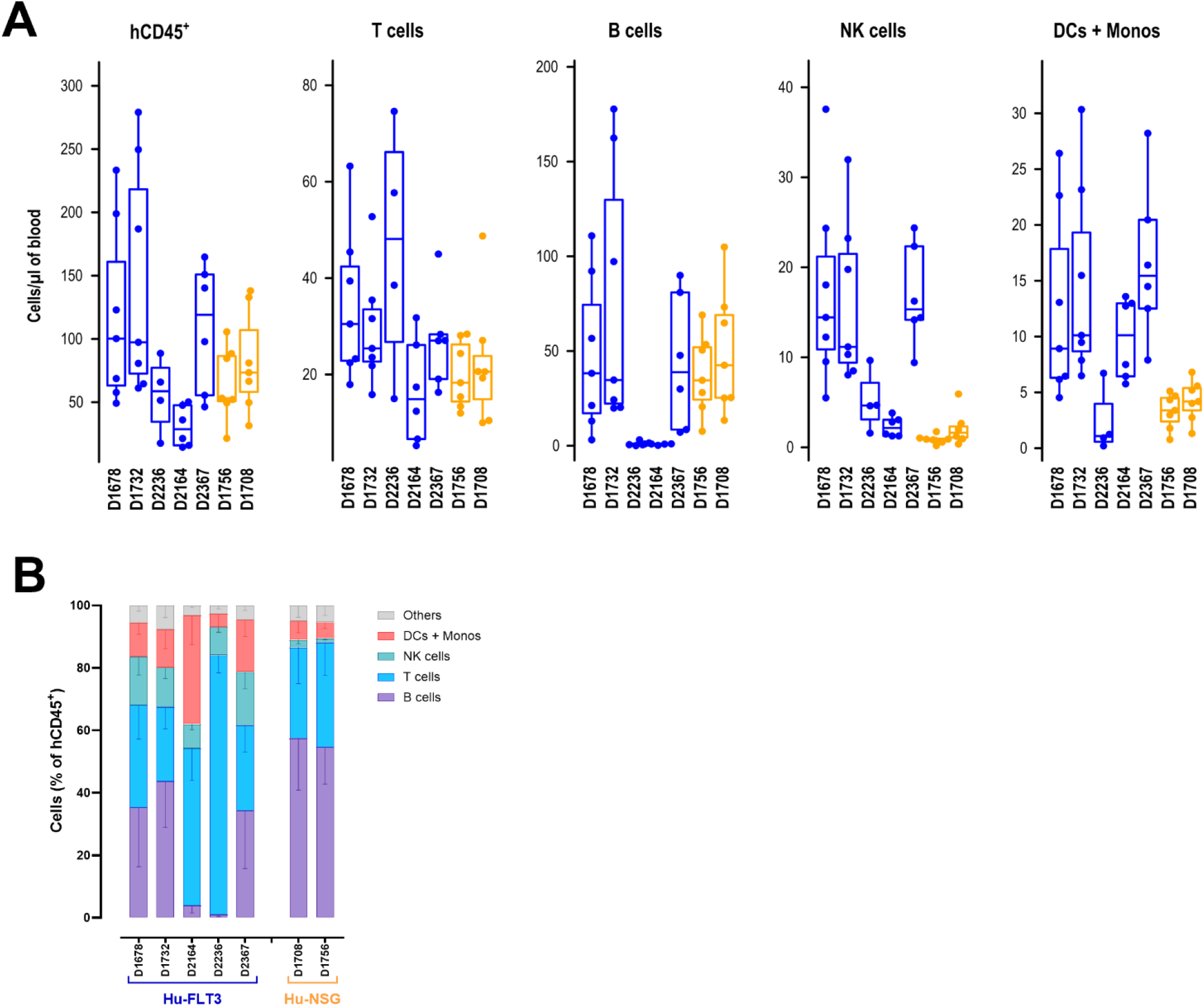
Donor-dependent variation of human hematopoietic engraftment in Hu-FLT3 and Hu-NSG mice at the earliest time point prior to infection, between week 15 and week 19 post-transplant. **A**. Frequencies of B cells (purple), T cells (blue), NK cells (green), and combined DCs/monocytes (red) in peripheral blood. Hu-FLT3 mice were humanized with CD34^+^ cells from five donors and Hu-NSG mice were humanized with CD34^+^ cells from two donors. **B**. Absolute numbers of human CD45^+^ cells, T cells, B cells, NK cells, and combined DCs/monocytes in peripheral blood are shown for individual donors contributing to Hu-FLT3 (blue) and Hu-NSG (orange) cohorts. Each data point represents an individual animal, displayed as a box-and-whisker plot (median and interquartile range).

**Figure S3.**
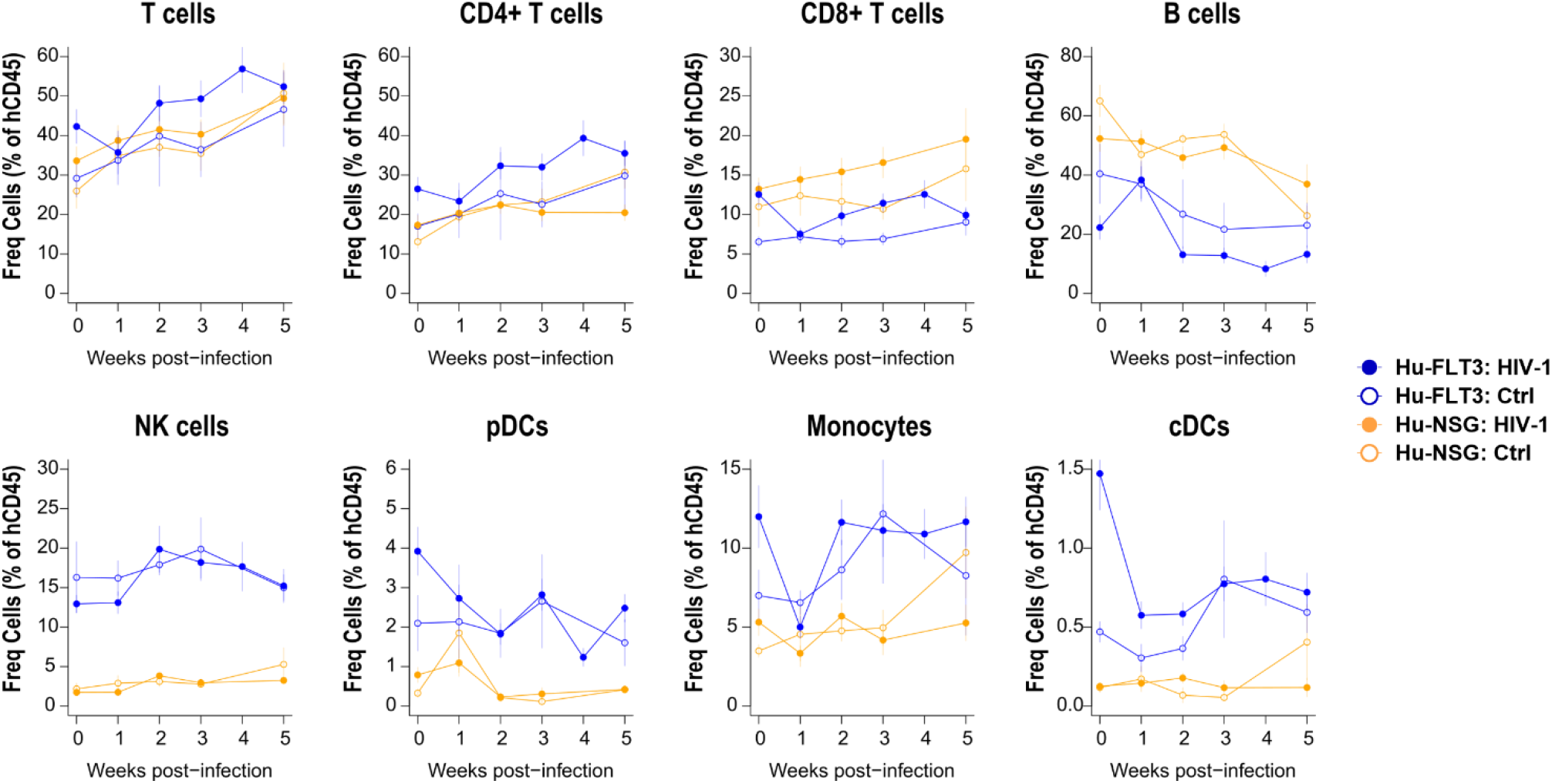
Immune cell dynamics in peripheral blood of Hu-FLT3 and Hu-NSG mice during the first five weeks of HIV-1 infection. Longitudinal frequencies of major human immune cell subsets (% of hCD45^+^) at 0, 1, 2, 3, and 5 wpi in Hu-FLT3 and Hu-NSG mice. Quantified populations include T cells, B cells, NK cells, pDCs, monocytes, cDCs, CD4^+^ T cells, and CD8^+^ T cells. Data are shown as mean ± SEM. Groups compared include Hu-FLT3 HIV-1 (n = 10), Hu-FLT3 Ctrl (n = 4, non-infected control group), Hu-NSG HIV-1 (n = 10), and Hu-NSG Ctrl (n = 4, non-infected control group).

**Figure S4.**
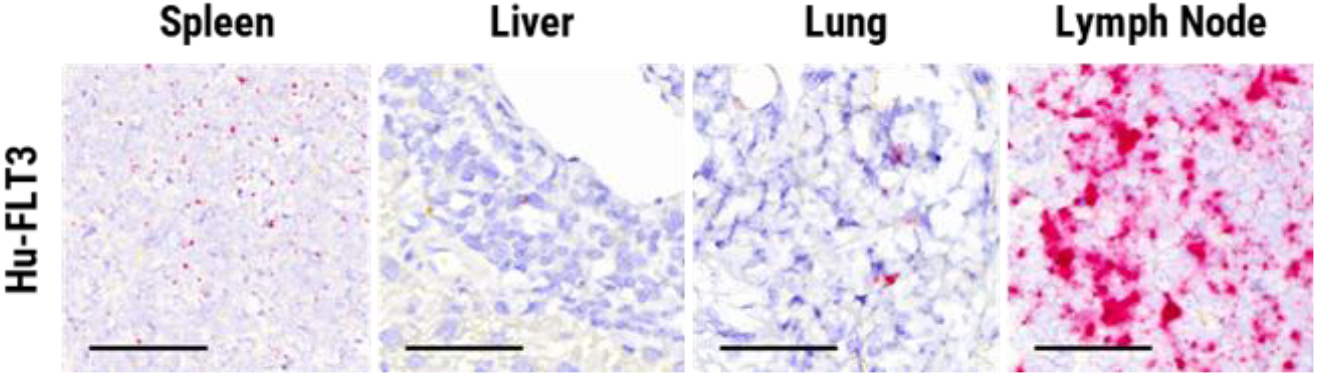
Detection of HIV-1 RNA in tissues from Hu-FLT3 mice at 5 wpi using RNAscope *in situ* hybridization. Representative images from spleen, liver, lung, and tracheobronchial lymph node are shown. HIV-1 RNA signal is shown in pink. Scale bar = 50 µm.

**Figure S5.**
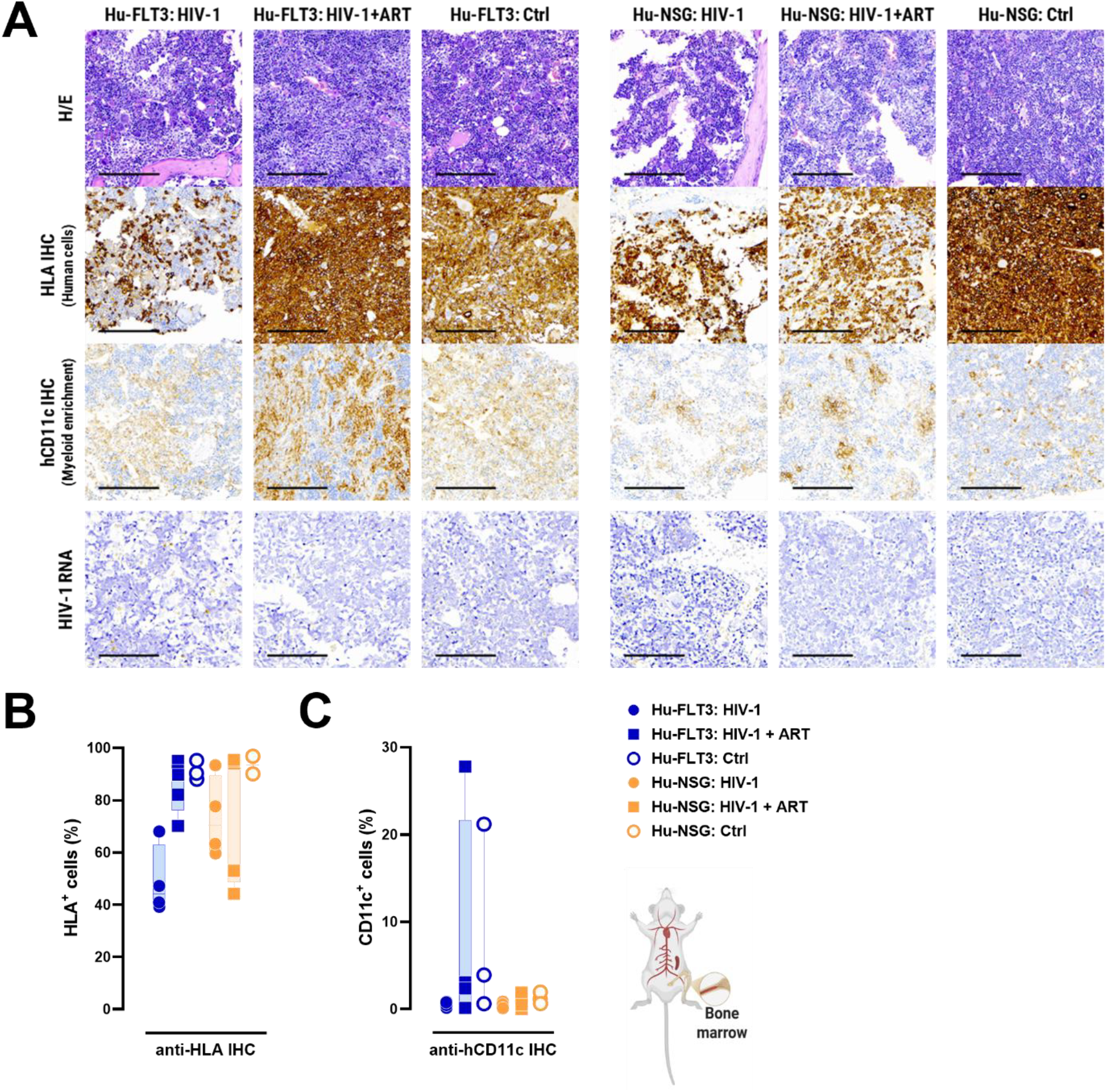
Human immune cell reconstitution and tissue-associated HIV-1 RNA in bone marrow of humanized mice. Hu-FLT3 and Hu-NSG mice were analyzed at the terminal timepoint (17 wpi) to compare only HIV-1 infected mice (Hu-FLT3, blue circles, n = 4; Hu-NSG, orange circles, n = 4;), HIV-1 infected on ART (Hu-FLT3, blue squares, n = 5; Hu-NSG, orange squares, n = 5), and non-infected controls (Hu-FLT3 n = 3; Hu-NSG n = 3; open circles). Bone marrow sections were evaluated by H/E staining, HLA and CD11c immunohistochemistry, and RNAscope HIV-1 RNA *in situ* hybridization. Quantification was performed in QuPath software as percent positive cells for HLA and CD11c within tissue. HIV-1 RNA was undetectable in bone marrow and was not quantified. **A**. Representative histological images of bone marrow sections from Hu-FLT3 and Hu-NSG mice stained with indicated cellular markers (brown) and HIV-1 RNA (pink). Scale bar = 200 µm. **B**. Quantification of HLA-positive cells. **C**. Quantification of human CD11c-positive cells. In **B** and **C**, individual mice are shown as box-and-whisker plots.

